# EpiMethylTag simultaneously detects ATAC-seq or ChIP-seq signals with DNA methylation

**DOI:** 10.1101/549550

**Authors:** Priscillia Lhoumaud, Gunjan Sethia, Franco Izzo, Theodore Sakellaropoulos, Valentina Snetkova, Simon Vidal, Sana Badri, MacIntosh Cornwell, Dafne Campigli Di Giammartino, Kyu-Tae Kim, Effie Apostolou, Matthias Stadtfeld, Dan Avi Landau, Jane Skok

**Affiliations:** New York University Langone Health, New York, NY, USA; New York Genome Center, New York, NY, USA; Meyer Cancer Center, Weill Cornell Medicine, New York, NY, USA; Laura and Isaac Perlmutter Cancer Center, NYU School of Medicine, New York, NY 10016, USA; Skirball Institute of Biomolecular Medicine, Department of Cell Biology, Helen L. and Martin S. Kimmel Center for Biology and Medicine, Laura and Isaac Perlmutter Cancer Center, New York, NY, USA; Sanford I. Weill Department of Medicine, Sandra and Edward Meyer Cancer Center, Weill Cornell Medicine, New York, NY, USA; Institute of Computational Biomedicine, Weill Cornell Medicine, New York, NY, USA

**Author notes:** Email addresses.

**Keywords:** DNA methylation, chromatin accessibility, ChIP, ATAC, CTCF, KLF4

## Abstract

Activation of regulatory elements is thought to be inversely correlated with DNA methylation levels. However, it is difficult to determine whether DNA methylation is compatible with chromatin accessibility or transcription factor (TF) binding if assays are performed separately. We developed a fast, low input, low sequencing depth method, EpiMethylTag that combines ATAC-seq or ChIP-seq (M-ATAC or M-ChIP) with bisulfite conversion, to simultaneously examine accessibility/TF binding and methylation on the same DNA. Here we demonstrate that EpiMethylTag can be used to study the functional interplay between chromatin accessibility and TF binding (CTCF and KLF4) at methylated sites.

## Introduction

The role of DNA methylation (DNAme) in gene regulation has been widely described [1-4]. In general, methylation is thought to reduce accessibility and prohibit TF binding at enhancers and promoters [5, 6]. Nevertheless, TFs are also known to bind methylated DNA [2], but due to limitations in the techniques available for this kind of analysis, few genome wide studies have been performed. As a result, we still know very little about the DNA sequence and chromatin context of TF binding at methylated sites and its significance to gene regulation.

Several techniques have been developed to measure DNAme, some more comprehensive than others. Whole genome bisulfite sequencing (WGBS) covers all genomic regions, however, to achieve sufficient sequencing coverage is costly. The alternative, reduced representation bisulfite sequencing (RRBS), which requires less sequencing depth, preferentially captures CpG-dense sequences known as CpG islands that can potentially act as regulatory elements [7]. Nevertheless, both techniques require additional assays on different batches of cells to elucidate the interplay between DNAme, DNA accessibility and TF binding and this does not satisfactorily address the issue of compatibility. Current techniques that simultaneously analyze methylation together with TF binding or accessibility (NOME-seq [8], HT-SELEX [9], ChIP-bisulfite [10], BisChIP-seq [11], ChIP-BisSeq [12]) have drawbacks such as analysis of DNA rather than chromatin and the requirement of large amounts of input DNA or high sequencing costs.

To circumvent the high input and sequencing expenses associated with WGBS and existing ChIP combined with bisulfite conversion protocols [10-12], we developed ‘EpiMethylTag’. This technique combines ATAC-seq or ChIPmentation [13, 14] with bisulfite conversion (M-ATAC or M-ChIP, respectively) to specifically determine the methylation status of accessible or TF-bound regions in a chromatin context. EpiMethylTag is based on an approach that was originally developed for tagmentation-based WGBS [15, 16]. It involves use of the Tn5 transposase, loaded with adapters harboring cytosine methylation (**Table S1**).

For M-ATAC or M-ChIP, tagmentation occurs respectively on nuclear lysates as per the conventional ATAC-seq protocol [13], or during chromatin immunoprecipitation as per the ChIPmentation protocol [14]. Following DNA purification, the sample is bisulfite converted and PCR amplified for downstream sequencing (**Fig. 1a**). As shown in **Fig. 1a**, EpiMethylTag can determine whether DNAme and accessibility/TF binding are mutually exclusive (scenario 1) or can coexist in certain locations (scenario 2). The protocol requires lower levels of immunoprecipitated DNA, less sequencing depth, is quicker than existing methods and can be analyzed using a pipeline we developed that is publicly available online on Github (“https://github.com/skoklab/EpiMethylTag”).

**Fig. 1.**
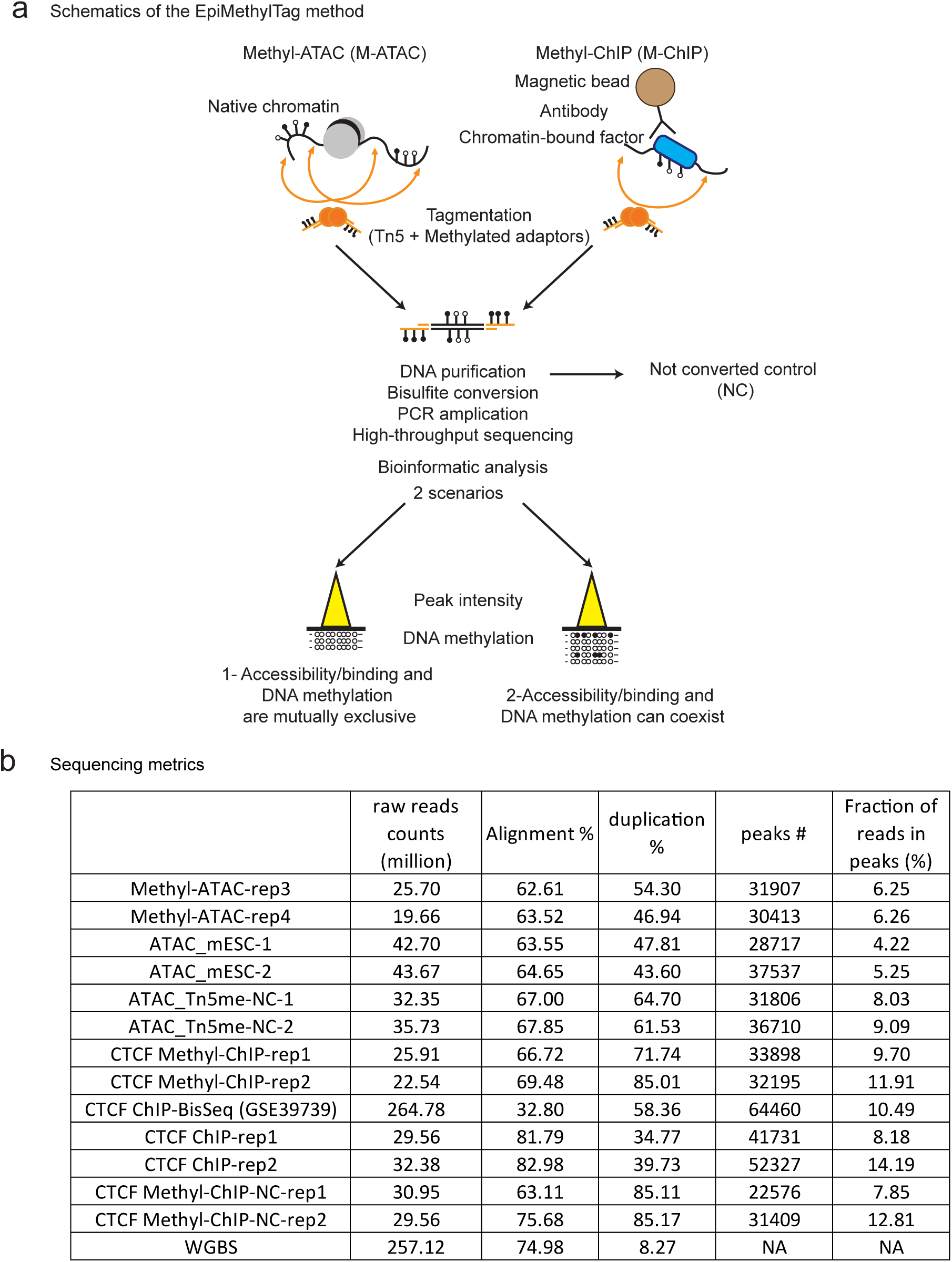
EpiMethylTag is a reproducible method to test whether DNAme can coexist with TF binding (CTCF) or chromatin accessibility. **a** Schematic overview of the EpiMethyTag method showing two possible outcomes. **b** Sequencing metrics indicating the total number of reads in million, the alignment and duplication percentages, the number of peaks and the fraction of reads in peaks (in percentage) for each sample as compared to public data (CTCF ChIP-BisSeq and WGBS).

## Results

### EpiMethylTag is a reproducible method for testing the compatibility of DNAme with TF binding or chromatin accessibility

M-ATAC and CTCF M-ChIP were performed in duplicate on murine embryonic stem cells (mESC). As controls, we collected aliquots before bisulfite conversion, ATAC-seq and CTCF ChIPmentation with Nextera transposase [13, 14]. Sequencing metrics are shown in **Fig. 1b** and **Table S2**. The price is around 10 times lower than WGBS given that fewer reads are necessary. As shown in **Fig. 2a and 2b**, genome coverage was highly reproducible between M-ATAC replicates and highly correlated with regular ATAC-seq and M-ATAC signal before bisulfite treatment. Thus, bisulfite treatment, or the use of a different transposase does not result in signal bias. High reproducibility was also seen for CTCF M-ChIP, and we observed consistency between our results and data generated by CTCF ChIP-BisSeq, a similar technique that was performed using 100ng of immunoprecipitated DNA (as opposed to less than 1ng using our method) and sequenced more deeply at higher cost [12] (**Fig. 2a and 2b, Table S2**). Of note, bisulfite conversion does not affect the number of peaks detected, the Jaccard index of peak overlap (**Figure S1a-b**) or the signal within peaks (**Figure S1c**, Pearson correlations above 0.8), although it leads to shorter reads (**Figure S2**). Of note, average methylation was higher at the edges of the peaks than at the midpoint (**Figure S3**). Comparable DNA methylation levels were found in M-ATAC and CTCF M-ChIP replicates, Pearson correlation = 0.76 and 0.84, respectively (**Figure S4a and S4b**).

**Fig. 2.**
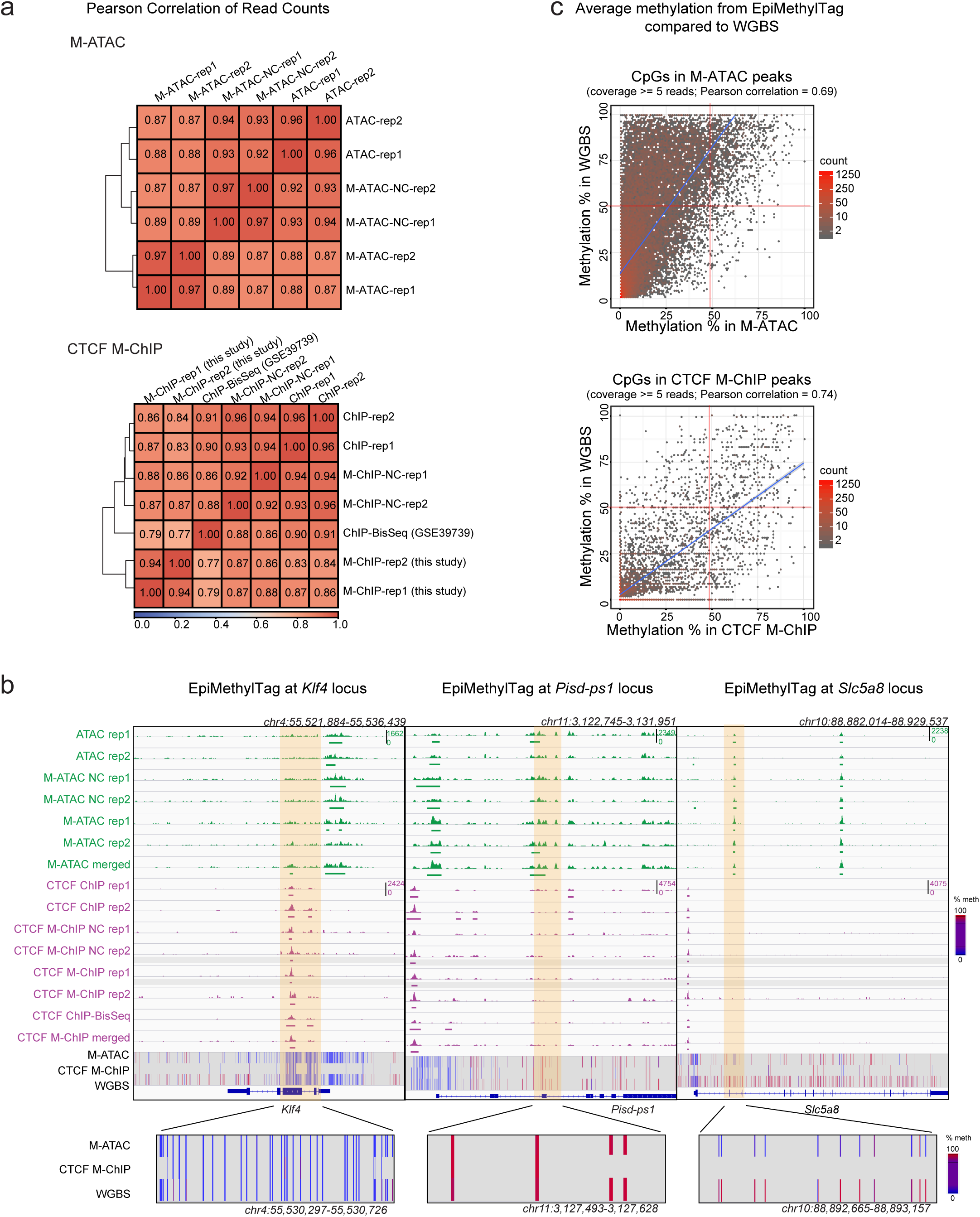
EpiMethylTag is a reproducible method for testing whether DNAme can coexist with TF binding (CTCF) or chromatin accessibility genome-wide. **a** Pearson correlation of read counts comparing M-ATAC with unconverted samples (NC) and regular ATAC-seq (top), and CTCF M-ChIP with unconverted samples, a sample from the Schubeler lab generated using ChIP-BisSeq [1] (GSE39739) and regular CTCF ChIP-seq (bottom). **b** Representative IGV screenshots of EpiMethylTag, at the *Klf4* locus (left panel), the *Pisd-ps1* locus (middle panel), and the *Slc5a8* locus (right panel). ATAC and M-ATAC in green, CTCF in purple and DNA methylation from merged M-ATAC, merged CTCF M-ChIP and WGBS (methylation from 0% in blue to 100% in red). A zoom-in of methylation at the highlighted region is shown at the bottom of each example. The *Klf4* locus illustrates a region that has low methylation as detected by M-ATAC, CTCF M-ChIP and WGBS. The *Pisd-ps1* locus illustrates a region that has high methylation as detected by M-ATAC, CTCF M-ChIP and WGBS. The *Slc5a8* locus illustrates a region that has low methylation as detected by M-ATAC and high methylation as detected by WGBS. **c** Density plots of methylation from EpiMethyltag compared with WGBS. Only CpGs inside peaks and with at least 5 reads were considered. Top: average methylation of CpGs per M-ATAC peak in M-ATAC versus WGBS (Pearson Correlation = 0.69, p-value < 2.2e-16; bottom left corner: 27977 peaks, top left corner: 8408 peaks, top right corner: 1019 peaks, bottom right corner: 113 peaks). Bottom: average methylation per CTCF M-ChIP peak of CpGs in CTCF M-ChIP versus WGBS (Pearson Correlation = 0.74, p-value < 2.2e-16; bottom left corner: 6549 peaks, top left corner: 198 peaks, top right corner: 304 peaks, bottom right corner: 310 peaks).

In order to get higher coverage for subsequent DNA methylation analysis, peaks were called from merged M-ATAC and M-ChIP replicates and we focused our analysis only at CpGs within those peak regions covered by at least five reads, as methylation outside of M-ATAC and M-ChIP peaks has low coverage and is less reliable. We observe positive correlations between DNA methylation from WGBS and M-ATAC (**Fig. 2c**, top panel, Pearson correlation=0.69), and between methylation levels in M-ChIP and WGBS (**Fig. 2c**, bottom panel, Pearson correlation = 0.74). Similar results were observed with the previously published CTCF ChIP-BisSeq method [12] (GSE39739) (Pearson correlation = 0.83, **Figure S4c**) and when taking peaks that overlap between duplicates (**Fig. S4d-e**). In **Fig. 2b**, we highlight the *Klf4* gene, which harbors a peak of chromatin accessibility in the promoter and CTCF binding in the intragenic region associated with low methylation from both EpiMethylTag and WGBS assays (left panel). In contrast, the *Pisd-ps1* intragenic region contains accessible chromatin that coexists with high levels of DNA methylation as detected by both M-ATAC and WGBS (**Fig. 2b**, middle panel). Of note, the methylation observed comes from a bedGraph file, output from Bismark (see method section for details), which does not filter for cytosines with low read coverage. Therefore, high methylation observed in CTCF M-ChIP may not be reliable as this region harbors a weak CTCF signal with a low read coverage (**Table S3**). Interestingly, a proportion of M-ATAC peaks exhibited an intermediate-to-high average methylation level in deeply sequenced WGBS [17], but low methylation in M-ATAC (**Fig. 2c**, top panel, top left corner) as illustrated at the *Slc5a8* locus (**Fig. 2b**, right panel, **Table S3**). The peak highlighted within the *Slc5a8* locus harbors an average methylation of 18.685% for M-ATAC and 85.041% for WGBS. These data suggest that as expected open regions are less methylated than closed regions within a population of cells, but that accessibility and methylation can coexist at a small subset of genomic locations, which are depleted for promoter regions and associated with low transcription (**Figure S4f-g**). Importantly, M-ATAC is able to identify methylation levels within ATAC peaks, information that cannot be retrieved integrating data from separate WGBS and ATAC-seq experiments.

### M-ATAC reveals a complex interplay between accessible chromatin and DNA methylation

For further analysis, we separated CpGs in M-ATAC peaks according to percentage of methylation (low 0-20%, intermediate 20-80% and high >80%) and read coverage (high > 50 reads and low 5-50 reads) as follows: #1: Low methylation/High coverage (22932 CpGs); #2: Low Methylation/Low coverage (1348931 CpGs); #3: Intermediate methylation/Low coverage (39321 CpGs); #4: High methylation/Low coverage (1652 CpGs) (**Fig. 3a**). As expected, coverage and methylation from M-ATAC are anticorrelated and we did not detect any CpGs with intermediate or high methylation with high ATAC coverage (>50 reads). A similar pattern was observed while taking only CpGs presents in peaks that overlap between M-ATAC replicates (**Figure S5a**). Of note, this pattern was not detected in WGBS where a more stable coverage is observed independent of methylation levels resulting in only 3 groups (**Figure S5b**) as opposed to the 4 groups seen with methyl-ATAC (**Fig. 3a**). CpGs in low methylation M-ATAC groups 1 and 2 were enriched at promoters, while CpGs in intermediate and high methylation M-ATAC groups 3 and 4, were enriched in intragenic and intergenic regions, as compared to the full set of M-ATAC peaks (**Fig. 3b**). The average methylation was more negatively correlated with transcriptional output for CpGs at promoters (**Fig. 3c**) than for intragenic CpGs (**Figure S5c**). Heatmaps for M-ATAC read coverage intensity highlight the reproducibility of signal between individual replicates. Merged replicates were used for downstream analysis (**Figure S5d**). Intriguingly, H3K4me1 showed a pronounced enrichment at CpGs with high levels of methylation (group 4) at promoter regions (**Fig. 3d and Figure S5e**). In contrast, H3K27ac and H3K4me3 were enriched at CpGs with low levels of methylation (groups 1 and 2), for both promoters and non-promoters.

**Fig. 3.**
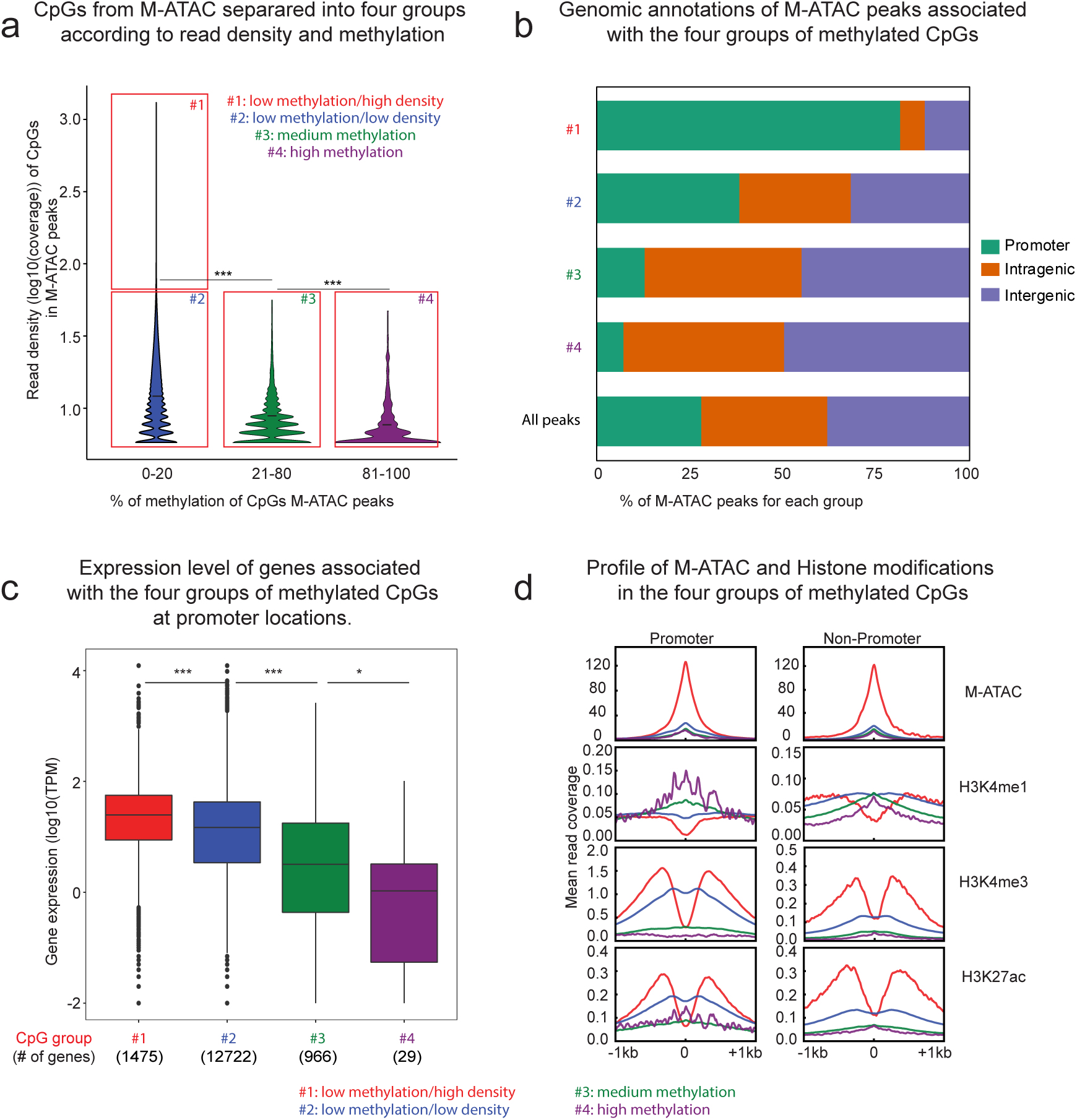
M-ATAC reveals a complex interplay between accessible chromatin and DNA methylation. **a** CpGs in M-ATAC peaks from merged replicates were divided into four groups according to methylation and coverage status: 1. Low Methylation (<20%) + High coverage (>50 reads) (22932 CpGs). 2. Low Methylation + Low coverage (5 to 50 reads) (1348931 CpGs). 3. Intermediate methylation (20-80) + Low coverage (5 to 50 reads) (39321 CpGs). 4. High methylation (>80%) + Low coverage (5 to 50 reads) (1652 CpGs). *** P<1e-300 between groups #1 + 2 and group #3, ***P=3.25e-109 between groups #3 and 4 (Wilcoxon text). **b** Genomic annotations for M-ATAC peaks corresponding to the 4 groups from **Fig. 3a** as well as the full list of M-ATAC peaks. Promoter: TSS - 3kb and +3kb; intragenic: introns, exons, 5’UTR, 3’UTR and TTS, intergenic: distal from promoter >1kb and non-coding RNAs. **c** Expression level of genes associated with the four groups of methylated CpGs from in **Fig. 3a**, for the CpGs at promoters. ***P=4.2e-33 between groups #1 and 2, ***P=2.8e-75 between groups #2 and 3, *P=0.034 between groups #3 and 4 (Wilcoxon test). **d** Average profile of M-ATAC, H3K4me1, H3K4me3 and H3K27ac signal associated with the four groups of methylated CpGs from **Fig. 3a** at promoters versus non-promoters. Of note, the small number of promoters in group 4 gives an unsmooth pattern for marks such as H3K4me1 and H3K27ac.

### CTCF M-ChIP enables analysis of DNA methylation of distinct CpGs in the CTCF motif

As a case study, CTCF M-ChIP was used to analyze the impact of DNAme on CTCF binding in M-ATAC peaks harboring a CTCF motif (**Fig. 4a**, top panel). M-ATAC groups 2 and 3 comprise the vast majority of CpGs, more CTCF peaks, motifs and a proportional higher number of CpGs within CTCF motifs (**Figure S5f**). However the percentage of CpGs within CTCF motifs in each group is fairly constant: between 1.26% to 1.93% of CpGs). Of note, *de novo* CTCF motifs in CTCF ChIP-seq and Methyl-ChIP peaks were comparable to the MA0139.1 motif from the Jaspar database (**Figure S6a**). CTCF occupancy has been inversely correlated with DNA methylation [18]. This finding is consistent with our analyses (**Figure S6b-d**). Although CTCF peaks are associated with all levels of CpG methylation within CTCF motifs, as illustrated in **Figure S6e**, the majority of CTCF peaks harbor reduced methylation (**Figure S6f**). In the context of CpGs in M-ATAC peaks, our data also demonstrates that the CTCF motif has an enriched CTCF intensity at CpGs with low and intermediate levels of methylation (groups 2 and 3) compared to CpGs with low and high levels of methylation (groups 1 and 4) (**Fig. 4a**, bottom panel). The highest binding is found in groups 2 and 3, compared to groups 1 and 4 that harbor reduced CTCF enrichment. Group 2 displays a wide range of accessibility (**Figure S5d-e**), with the most open regions of group 2 resembling group 1, and the most closed regions of this group being similar to that of group 3. Interestingly, even though there are more CpGs in CTCF motifs in group 1 compared to group 4 (**Figure S5f**, 288 versus 25 CpGs), group 1 shows a lower level of CTCF enrichment than group 4. This may be due to the confidence of attributing CpGs to a specific group. As shown in **Figure S6g**, for all clusters, more than half of the CpGs have a high probability of being in the assigned group (> 72%). These data provide insight into CTCF binding and suggest an anticorrelation between high accessibility and high methylation.

**Fig. 4.**
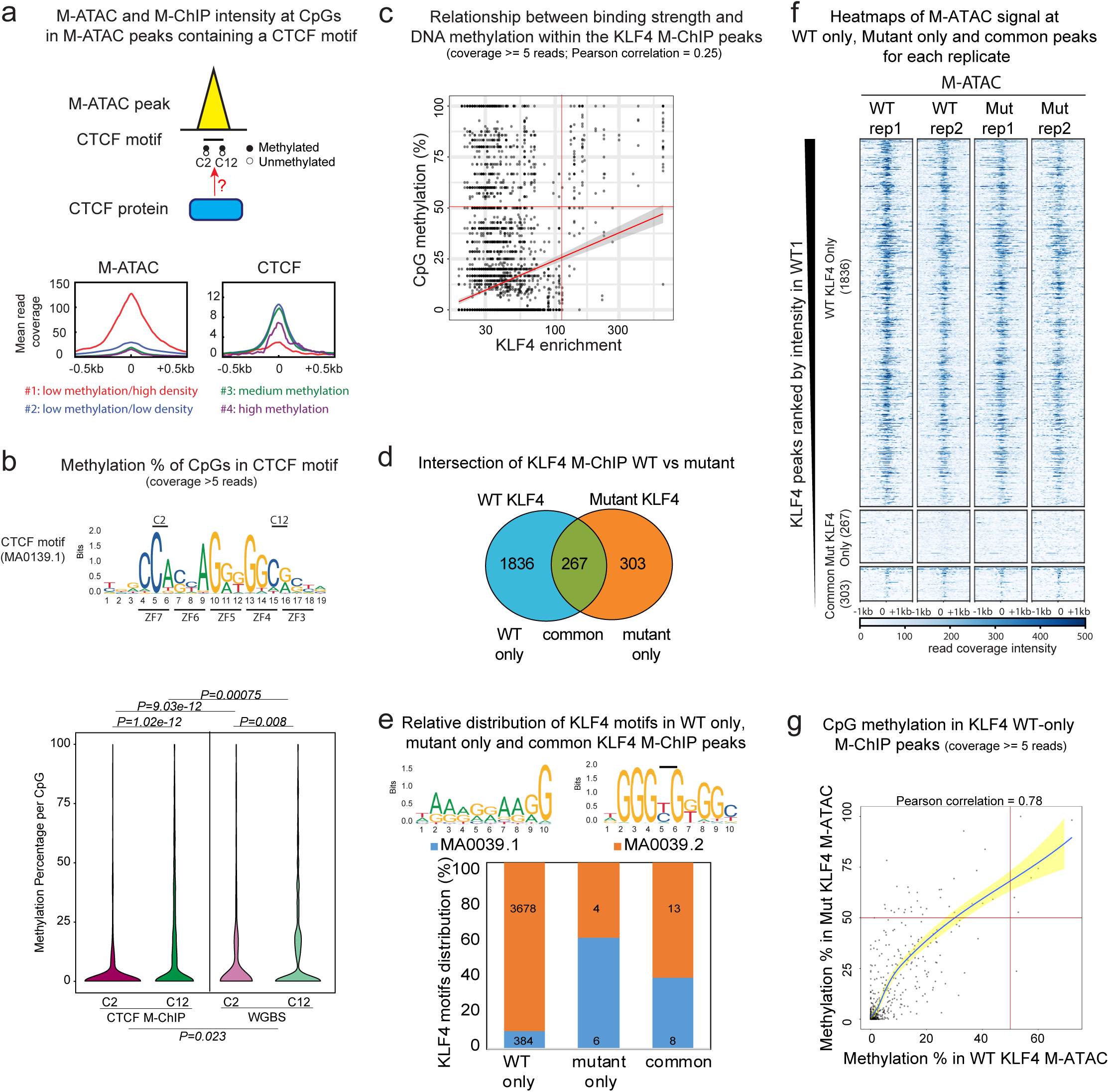
M-ChIP enables analysis of DNA methylation binding by CTCF and KLF4. **a** Top: Schematic illustration representing an ATAC-seq peak with a CTCF motif and CTCF occupancy dependent on C2 and C12 methylation. Bottom: average profiles of M-ATAC (left) and CTCF M-ChIP (right) intensity at CpGs in a CTCF motif within M-ATAC peaks for the four groups of CpGs (group #1: 288 CpGs, group #2: 17133 CpGs, group #3 CpGs: 758, group #4: 25 CpGs). **b** top: CTCF motif from JASPAR database (MA0139.1). The 2 key CpG positions (C2 and C12) are indicated and bottom: violin plots of methylation percentage from CTCF M-ChIP and WGBS, at C2 and C12 positions into CTCF motif (MA0139.1). ***P=1.02e-12 for C2 CTCF M-ChIP versus C12 CTCF M-ChIP (Wilcoxon test), **P=0.008 for C2 WGBS versus C12 WGBS (Wilcoxon test), ***P=9e-12 for C2 CTCF M-ChIP versus C2 WGBS (Wilcoxon test, paired), ***P=0.00075 for C12 CTCF M-ChIP versus C12 WGBS (Wilcoxon test, paired), *P=0.023 for CTCF M-ChIP versus WGBS (logistic regression model). **c** Scatter plot showing the relationship between binding strength and CpG methylation within the KLF4 M-ChIP peaks (Pearson Correlation = 0.25; bottom left corner: 5138 CpGs, top left corner: 578 CpGs, top right corner: 104 CpGs, bottom right corner: 60 CpGs). **d** Venn diagram showing the overlap between WT and mutant KLF4 M-ChIP peaks. **e** Top: Illustration of KLF4 motifs from the Jaspar database (MA0039.1 and MA0039.2). The black bar represents the potential CpGs present in the MA0039.2 motif. Bottom: histogram showing the relative distribution of KLF4 motifs in WT, mutant and common KLF4 M-ChIP peaks using FIMO from the MEME suite. Absolute numbers of each motif are indicated. **f** Heatmap showing M-ATAC signal intensity at KLF4 M-ChIP peaks that are specific to WT (1836 peaks), mutant (267 peaks) or common between both conditions (303 peaks). **g** Average cytosine methylation from M-ATAC in WT versus mutant KLF4 expressing cells in WT specific KLF4 M-ChIP peaks (Pearson Correlation = 0.78, p-value < 2.2e-16).

The MA0139.1 CTCF motif incorporates 2 CpGs: C2 and/or C12 (**Fig. 4b**, top panel). According to the CTCF logo, we identified more CpGs at position C12 than C2 in the CTCF M-ChIP peaks (4884 versus 921 CpGs, respectively, considering only the CpGs covered by at least 5 reads in both M-ChIP and WGBS). Consistent with the findings from a recent study that analyzed CTCF binding using oligonucleotides rather than genomic DNA [19], CTCF M-ChIP detected higher levels of methylation at C12 compared to C2 (**Fig. 4b**, bottom panel, compare CTCF M-ChIP C2 versus C12, p-value = 1.02e-12). Importantly, CTCF M-ChIP is more suitable than WGBS for detecting the differences (**Fig. 4b**, bottom panel, compare CTCF M-ChIP versus WGBS, p-value = 0.023). In addition, we found that bi-methylation at both CpGs within the same read is slightly enriched compared to what is expected by random chance (0.97% versus 0.05%) (**Figure S7a**, = χ^2^1531, p-value < 0.001). CTCF signal intensity is relatively comparable at the 4 combinations of methylation, with a slight increase for C2 being methylated and C12 unmethylated (**Figure S7b**), however the biological significance of this remains to be determined. Nonetheless, sequence variation at the C2 and C12 positions appears to have no effect on methylation levels (**Figure S7c**).

### KLF4 M-ChIP enables characterization of WT versus mutant KLF4 R462A binding

Pioneer transcription factors need to access target genes that are inaccessible and whose enhancer and promoter sequences may be methylated. A recent study has shown that a minority of transcription factors (47 out of 1300 examined) including KLF4 can bind to methylated CpG sites [2]. A scatter plot of KLF4 M-ChIP in WT mESCs shows that the majority of CpGs in KLF4 peaks display low peak intensity and low methylation (**Fig. 4c**). However, in contrast to CTCF, the small fraction of peaks with the highest peak intensity also display the highest methylation levels. The study mentioned above [2], revealed that distinct zinc fingers on KLF4 mediate KLF4’s binding activity with methylated and unmethylated DNA. Residue arginine 458 on human KLF4 was shown to be important for binding to the methylated motif CCmCpGCC [2] (similar to the Jaspar motif MA0039.2 for mouse KLF4). In the mouse protein the equivalent arginine residue lies at position 462.

In order to investigate the binding of KLF4 to methylated DNA, we used *Klf4*^-/-^ mESCs [20] that express either a WT or mutant version of KLF4 in which arginine 462 has been replaced by alanine (R462A) **(Figure S8a-b)**. We performed KLF4 M-ChIP in both WT and mutant expressing mESC in duplicates. Intersections between replicates were used to identify peaks specific to (i) WT or (ii) mutant versions of KLF4 and (iii) those that were common to both (**Fig. 4d**). Heatmaps confirm the binding specificity of the two versions of KLF4 and reveal the high reproducibility between duplicates (**Figure S8c**).

We searched for mouse KLF4 motifs from the Jaspar database, using the FIMO tool from the MEME suite. The two motifs that were identified, MA0039.2 and MA0039.1 can be distinguished by the presence and absence of a CpG dinucleotide, respectively (**Fig. 4e**, top). The wild-type version of KLF4 has a strong preference for motif MA0039.2 while the mutant loses this preference. Overall the mutant protein has reduced binding to both motifs (**Fig. 4e**, bottom).

Because of the low numbers of consensus KLF4 motifs in common and KLF4 mutant specific peaks, we decided to focus our downstream analysis only on WT specific peaks. M-ATAC experiments conducted in duplicates in both WT and Mutant KLF4 expressing cells show that KLF4 peaks present only in the WT condition are accessible, while Mutant only KLF4 peaks are found at inaccessible sites (**Fig. 4f**). This result together with the motif findings (**Fig. 4e**) suggest that the Mutant KLF4 binding alone occurs at inaccessible sites where there is no consensus KLF4 motif. Thus, this mutation abrogates binding at consensus KLF4 motifs. The functional significance of binding of Mutant KLF4 at ectopic sites remains to be investigated. WT specific KLF4 peaks harbor similar DNA accessibility in both WT and Mutant conditions so it is not clear why the Mutant protein does not bind. To investigate, we analyzed DNA methylation at these sites using M-ATAC, M-ChIP and public WGBS from WT mESCs. The levels of methylation obtained from M-ATAC were also compared for cells expressing WT and Mutant KLF4 within the WT specific KLF4 M-ChIP peaks. In the scatter plots shown in **Fig. 4g** and **Figure S8d**, most of the CpGs display low levels of methylation in any condition (bottom left corner). Thus, methylation levels do not explain the absence of Mutant KLF4 binding at these sites.

## Discussion

We developed a new method, “EpiMethylTag”, that allows the simultaneous analysis of DNA methylation with ChIP-seq or ATAC-seq. EpiMethylTag can be used to analyze the methylation status and coincident accessibility or binding of other chromatin bound transcription factors. Importantly our approach is a fast, low input, low sequencing depth method that can be used for smaller cell populations than existing methods and can adapted for rare cell populations. Specifically, our M-ChIP protocol significantly reduces the input for DNA binding factors such as CTCF. The only published genome-wide ChIP-Bis-Seq for CTCF [12] used 100 ng of immunoprecipitated DNA. Using a Tn5 transposase successfully allowed us to use less than 1ng of immunoprecipitated DNA followed by bisulfite conversion. The number of cells required to obtain 1ng of ChIPped DNA will vary depending on the protocol and the antibody used. ChIP-bisulfite [10] and BisChIP-seq [11] use lower cell numbers for H3K27me3. However, such histone modifications in general require less cells for ChIP on TFs such as CTCF or KLF4 because they cover a higher portion of the genome. Although it has not been tested, our protocol may also lower the number of cells required for M-ChIP of histone modifications.

EpiMethylTag confirmed that as a general rule, DNA methylation rarely coexists with DNA accessibility or TF binding. Nonetheless, we found M-ATAC peaks of low signal intensity that overlapped with DNA methylation. These peaks were located predominantly in intragenic and intergenic regions and associated with low transcriptional output at gene promoters. This data identifies a class of promoters with high accessibility, high levels of methylation, high H3K4me1, low K3K4me3 and low H3K27ac (**Fig. 3d**). The biological relevance of such ‘poised promoters’, remains to be determined.

Of note, a recent publication used the same design for the Methyl-ATAC aspect of EpiMethylTag method [21]. As with our approach they show that mATAC-seq detects methylation patterns that agree with both WGBS and Omni-ATAC (improved normal ATAC-seq [22]). By comparing parental and DNMT1 and DNMT3B double knockout HCT116 cells they identified ATAC peaks with increased accessibility that were bound by TFs only in the demethylated cells. However, they did not adapt their approach to analysis of methylated ChIP-seq peaks as we have done. Here we used M-ChIP to characterize the binding of both CTCF and KLF4 to motifs in the context of DNA methylation.

Methylation within CTCF motifs is known to be anticorrelated with CTCF binding [3]. Our analysis revealed that M-ATAC peaks containing a CTCF motif have an enriched CTCF intensity at CpGs with intermediate levels of methylation as opposed to low and high levels of methylation. In addition, CTCF M-ChIP revealed that methylation at CpG C2 is lower than at CpG at position C12, a finding that suggests methylation at C2 could have a stronger negative impact on CTCF binding than methylation at C12. Differences of this sort could not be detected by integrating CTCF ChIP-seq with WGBS (**Fig. 4b**).

We further demonstrate that M-ChIP could be used to characterize the profiles and methylation status of common WT and mutant KLF4 R462A binding sites. Thus, methylation levels do not explain the absence of Mutant KLF4 binding at these sites and it appears that the mutant does not bind the consensus motif so we cannot investigate the relationship between methylation in the KLF4 motif and binding of WT versus Mutant KLF4 (**Fig. 4f-g**). While the biological significance of such differences remains to be investigated, our data demonstrate that EpiMethylTag can be used to provide information about the methylation status of the binding sites for the WT and mutant proteins. This information could not be obtained by performing separate methylation and ChIP-seq experiments.

## Conclusion

In sum, M-ATAC and CTCF M-ChIP reveal a complex interplay between accessible chromatin, DNA methylation and TF binding that could not be detected by WGBS. EpiMethylTag can be used to provide information about the DNA sequence and chromatin context of TF binding at methylated sites and its significance to gene regulation and biological processes. This technique can also be adapted for single cell analysis.

## Methods

### Cell culture

Mouse embryonic stem cells were provided by Matthias Stadtfeld. Briefly, KH2 embryonic stem cells (ESCs) [23] were cultured on irradiated feeder cells in KO-DMEM (Invitrogen) supplemented with L-glutamine, penicillin/streptomycin, nonessential amino acids, β-mercaptoethanol, 1,000 U/mL LIF, and 15% FBS (ESC medium). To remove feeder cells from ESCs, cells were trypsin digested and pre-plated in ESC medium for 30 min. Supernatant containing ESCs was used for further experiments.

### KLF4 expression

Mouse KLF4 has been cloned into pHAGE2-tetO-MCS-ires-tdTomato vector (obtained from Matthias Stadfeld’s lab, [24]) for the production of lentiviruses, using the following primers:

Fwd: 5’– gcggccgcATGGCTGTCAGCGACGCTCT

Rev: 5’– ggatccTTAAAAGTGCCTCTTCATGTGTAAGG

KLF4 R462A mutation has been generated using the site-directed mutagenesis kit from Agilent #210518. HEK 293T cells were used for the production of lentiviruses; obtained from ATCC (cat. No. CRL 3216). Lentiviral infection of KLF4 knock-out mESC [20] was performed by spin-infection and the cells were transferred to feeders and expanded with puromycin. After selection, KLF4 expression was induced with doxycycline (1ug/ml) for 2 days. Finally, the cells were pre-seeded (30 mins) to remove the feeders and the ES cells were processed as described in the “Cell culture” section. KLF4 protein expression has been checked by western blot using an antibody from Santa-Cruz (#sc-20691, now discontinued) and using H3 as a loading control (anti-H3, Abcam, ab1791).

### Assembly of the transposase

Tn5 transposase was assembled with methylated adaptors as per the T-WGBS protocol[16]. Ten microliters of each adapter with incorporated methylated cytosines (Tn5mC-Apt1 and Tn5mC1.1-A1block; 100 μM each; **Table S1**) were added to 80 μl of water and annealed in a thermomixer with the following program: 95 °C for 3 min, 70 °C for 3 min, 45 cycles of 30 s with a ramp at −1 °C per cycle to reach 26 °C. Fifty microliters of annealed adapters were incubated with 50 μl of hot glycerol and 10 μl of this mixture was incubated with 10 μl of Ez-Tn5 transposase (from the EZ-Tn5 insertion kit) at room temperature for 30 min to assemble the transposome.

### ATAC-seq and M-ATAC

ATAC-seq and M-ATAC were performed with 50 thousand mESCs as per the original ATAC-seq protocol [13]. Cells were washed in cold PBS and resuspended in 50 μl of cold lysis buffer (10 mM Tris-HCl, pH 7.4, 10 mM NaCl, 3mM MgCl_2_, 0.1 % IGEPAL CA-630). The tagmentation reaction was performed in 25 μl of TD buffer (Illumina Cat #FC-121-1030), 2.5 μl Transposase (either the Nextera transposase (ATAC-seq) or the transposase containing the methylated adaptors (M-ATAC, see section “assembly of the transposase” for details), and 22.5 μl of nuclease free H_2_O at 37°C for 30 min. Purified DNA (on column with the Qiagen Mini Elute kit) either bisulfite converted (M-ATAC, see section “Bisulfite conversion” for details) or directly amplified (ATAC-seq, see “Amplification of ATAC-seq and ChIP-seq libraries” for details).

### ChIPseq and M-ChIP

ChIP-seq and M-ChIP were performed on mESC as per the original ChIPmentation protocol [14]. Five microliters of CTCF antibody (Millipore 07-729) or 25 µl of KLF4 antibody (R&D AF3158) were combined to protein A (for CTCF) or G (for KLF4) magnetic beads and added to sonicated chromatin (from 200 to 700bp, checked on agarose gel) from 10 million mESC, for 3 to 6 hours rotating in the cold room. Beads were washed as per the original ChIPmentation protocol [14]: twice with TF-WBI (20 mM Tris-HCl/pH 7.4, 150 mM NaCl, 0.1% SDS, 1% Triton X-100, 2 mM EDTA), twice with TF-WBIII (250 mM LiCl, 1% Triton X-100, 0.7% DOC, and 10mM Tris -HCl, 1mM EDTA) and twice with cold Tris-Cl pH 8.0 to remove detergent, salts, and EDTA. During the second wash, the whole reaction was transfered to a new tube to decrease tagmentation of unspecific chromatin fragments sticking to the tube wall. Beads were resuspended in 25 µl of the tagmentation reaction mix (10 mM Tris pH 8.0, 5 mM MgCl2, and 10% v/v dimethylformamide) and tagmentation was performed for 1 min at 37°C with either 1 μl of the Nextera transposase (ChIP-seq) or the transposase containing the methylated adaptors (M-ChIP, see section “assembly of the transposase” for details). Then, beads were washed twice with TF-WBI (20 mM Tris-HCl/pH 7.4, 150 mM NaCl, 0.1% SDS, 1% Triton X -100, and 2mM EDTA) and twice with TET (0.2% Tween -20, 10 mM Tris-HCl/pH 8.0, 1 mM EDTA). During the last wash, the whole reaction was transfered to a new tube to decrease carry-over of tagmented unspecific fragments stuck to the tube wall. Chromatin was eluted and decrosslinked by 70μl of elution buffer (0.5% SDS, 300 mM NaCl, 5 mM EDTA, 10 mM Tris HCl pH 8.0) containing 20μg of proteinase K for 2 hours at 55 °C and overnight incubation at 65 °C. Eluted and purified DNA was either bisulfite converted (CTCF M-ChIP, see section “Bisulfite conversion” for details) or directly amplified (CTCF ChIP-seq, see “Amplification of ATAC-seq and ChIP-seq libraries” for details).

### Bisulfite conversion

Purified DNA was bisulfite converted following the T-WGBS protocol[16] with the EZ DNA methylation kit (Zymo). Oligonucleotide replacement was performed by incubating 9 μl of tagmented M-ATAC or M-ChIP purified DNA with 2 ng of phage lambda DNA as carrier, 2 μl of dNTP mix (2.5 mM each, 10 mM), 2 μl of 10× Ampligase buffer and 2 μl of replacement oligo (Tn5mC-ReplO1, 10 μM; Table S1) in a thermomixer with the following program: 50 °C for 1 min, 45°C for 10 min, ramp at −0.1 °C per second to reach 37 °C. 1 μl of T4 DNA polymerase and 2.5 μl of Ampligase were added and the gap repair reaction was performed at 37 °C for 30 min. DNA was purified using SPRI AMPure XP beads with a beads-to-sample ratio of 1.8:1 and eluted in 50 μl of H2O. 5 μl were kept as an unconverted control sample, and 45 μl was bisulfite converted using the EZ DNA methylation kit (Zymo). Briefly, the gap repair reaction was performed by adding 5 μl of M-dilution buffer and 15 min incubation at 37 °C, and bisulfite treatment was performed by adding 100 μl of liquid CT-conversion reagent in a thermomixer with the following program: 16 cycles of 95 °C for 15 sec followed by 50 °C for 1 hour. Converted DNA was purified on a column and amplified (see section “Amplification of M-ATAC and M-ChIP libraries” for details).

### Amplification of ATAC-seq and ChIP-seq libraries

Purified DNA (20 μl) was combined with 2.5 μl of each primer and 25 μl of NEB Next PCR master mix as per the original ATAC-seq protocol [13]. For ATAC-seq, DNA was amplified for 5 cycles and a monitored quantitative PCR was performed to determine the number of extra cycles needed not exceeding 12 cycles in total to limit the percentage of duplicated reads. DNA was purified on column with the Qiagen Mini Elute kit. For ChIP-seq, DNA was amplified as per the ChIPmentation protocol [14] in a thermomixer with the following program: 72 °C for 5 min; 98 °C for 30 s; 14 cycles of 98 °C for 10 s, 63 °C for 30 s and 72 °C 30 s; and a final elongation at 72 °C for 1 min. DNA was purified using SPRI AMPure XP beads with a beads-to-sample ratio of 1:1 and eluted in 20 μl of H2O.

### Amplification of M-ATAC and M-ChIP libraries

Purified converted DNA was amplified as per the original T-WGBS protocol [16]. Briefly, 10 μl of DNA was combined with 1.25 μl of each primer (25 μM each) and 12.5 μl of high-fidelity system KAPA HiFi uracil+ PCR master mix. DNA was amplified for 5 cycles and a monitored quantitative PCR was performed to determine the number of extra cycles needed, not exceeding 12 cycles in total to limit the percentage of duplicated reads.

### Sequencing of the libraries and data processing

For ATAC-seq, ChIP-seq, M-ATAC and M-ChIP, libraries were quantified using Kapa qPCR kit and sequenced using the HiSeq 2500 for paired-end 50 bp reads). ChIP-seq for histone modifications in mESC were downloaded from GEO (H3K4me1: GSM1000121, H3K27ac: GSM1000126, H3K4me3: GSM1000124). Data processing was performed as per the pipeline available on Github (link: “https://github.com/skoklab/EpiMethylTag”). Briefly, reads were trimmed using trim-galore/0.4.4, and aligned to the mm10 assembly of mouse genome using bowtie2 [25] for ChIP-seq and ATAC-seq, and using Bismark/0.18.1 (bowtie2) [26] for M-ChIP and M-ATAC to account for bisulfite conversion. Reads with quality < 30 and duplicates were removed using Samtools/1.3 [27]. Peaks were called using Macs/2.1.0 [28] with the following parameters: -- qvalue 0.01 --nomodel --shift 0 -B --call-summits. Narrow peaks were considered for further analysis. Bigwigs were generated from bam files with RPKM normalization using Deeptools [29] for visualization on IGV.

### Bioinformatic analysis of data

The distribution of fragment lengths was assessed with Deeptools/2.3.3 with option “-- maxFragmentLength 1000”, and Pearson correlations of reads counts with Deeptools/2.3.3 and default parameters. Heatmaps and average profiles were performed on merged bigwig files using Deeptools/2.3.3. Default parameters from Bismark/0.18.1 (Bowtie2) [26] were used to generate coverage files containing methylation information. Only cytosines in a CpG context were used for subsequent analysis. For **Fig. 3d** and **Figure S5b**, the plots were centered on CpGs in M-ATAC peaks from the different groups highlighted in **Fig. 3a**. For **Fig. 4a**, lists of CpGs were subsampled using BEDTools [30] to consider only the CpGs inside CTCF motifs, and the average plots were centered on those CpGs. Genomic annotations were performed using ChIPseeker [31]. CTCF motif locations in CTCF M-ChIP/ChIP and M-ATAC, and KLF4 motifs in M-ChIP peaks were determined using the FIMO tool from MEME [32], with the motif PWM from Jaspar database (MA0139.1 for CTCF and MA0039.1 and MA0039.2 for KLF4). PWM was manually modified to look at methylation frequency at different combinations of C2 and C12 dinucleotides of CTCF motif. Scripts are available on Github (link: “https://github.com/skoklab/EpiMethylTag”) for the reviewers upon request and publicly upon acceptance. In order to account for possible lack of specificity of the anti-KLF4 antibody, we filtered out ChIP-seq peaks present in *Klf4*^*-/-*^ cells. Peaks shared or specific to either WT or mutant KLF4 were identified using BEDTools [30]. For the ChIP enrichment versus CpG methylation plots, we plotted the peak score versus the beta values of the CpG probes within the peaks, using peaks called via MACS2 for CTCF (**Figure S6b**) and via PeaKDEck for KLF4 (**Fig. 4c**).

To quantify the probability of clustering CpG probes into low-, medium-, and highly methylated groups we assumed that beta values (ie the sampling mean) is normally distributed with mean the beta value (b) and variance (b(1-b))/((n-1)) where n is the total number of reads. This allows us to quantify the probability that each probe belongs to its designated cluster as *P*(*b* < *C*_*h*_) – *P*(*b* < *C*_*l*_) where *C*_*h*_ and *C*_*l*_ are the high and low thresholds of the cluster respectively. In the **Figure S6g**, the points and corresponding contours are colored based on their designated cluster. The x-axis is the beta value and the y-axis is the probability that beta lies within the cluster limits. For all clusters, more than half of the CpGs have a high probability of being in the assigned group (> 72%).

### Figure legends

**Figure S1**. Peak calling in EpiMethylTag, ATAC-seq and CTCF ChIP-seq. **a** Table showing number of peaks called for each sample, using MACS2. b Jaccard indexes of peak intersections between ATAC, M-ATAC, M-ATAC-NC samples (left panel) and CTCF ChIP-seq, CTCF M-ChIP and CTCF M-ChIP-NC samples (right panel). Jaccard Index = (Intersection / (sample 1 + sample 2 – Intersection)). c Scatter plots showing correlation of signal within peaks (union of any peak found in either condition). PCC = Pearson Correlation Curve. The x and y axes represent the log2fold of read counts.

**Figure S2**. Read lengths for all ATAC, M-ATAC, M-ATAC unconverted (M-ATAC-NC), CTCF ChIP-seq, CTCF M-ChIP and CTCF M-ChIP unconverted (CTCF M-ChIP-NC) samples.

**Figure S3**. Average methylation in M-ATAC peaks for CpGs with coverage of at least 5 reads, relative to the position of the CpGs in the peak.

**Figure S4**. Density plots of average methylation correlations for cytosines with coverage of at least 5 reads. **a** Average cytosine methylation from a M-ATAC replicate 1 versus replicate 2 in M-ATAC peaks (Pearson Correlation = 0.76, p-value < 2.2e-16). **b** CTCF M-ChIP replicate 1 versus replicate 2 in CTCF M-ChIP peaks (Pearson Correlation = 0.84, p-value < 2.2e-16). **c** CTCF ChIP-BisSeq (GSE39739) from Dirk Schubeler lab versus WGBS in CTCF ChIP-BisSeq peaks (Pearson Correlation = 0.83, p-value < 2.2e-16). **d** Average methylation of CpGs per M-ATAC peak in M-ATAC versus WGBS, only for M-ATAC peaks that overlap between both replicates (Pearson Correlation = 0.67, p-value < 2.2e-16). **e** Average methylation per CTCF M-ChIP peak of CpGs in CTCF M-ChIP versus WGBS only for M-ChIP peaks that overlap between both replicates (Pearson Correlation = 0.80, p-value < 2.2e-16). **f** Genomic annotations of peaks grouped according to their average methylation from WGBS and M-ATAC (relative to **Fig. 3c**, top panel). Promoter: TSS - 3kb to +3kb; intragenic: introns, exons, 5’UTR, 3’UTR and TTS, intergenic: distal from promoter >3kb. **g** Transcriptional output for the 4 groups of M-ATAC peaks according to their average methylation from WGBS and M-ATAC from **Figure S4d**, for the cytosines at promoters (left panel, see **Figure S4d**). ***P=1.25e-28 between groups #1 and 2, ^NS^P=0.19 between groups #2 and 3, ^NS^P=0.58 between groups #3 and 4 (Wilcoxon test), and for the cytosines at intragenic regions (right panel, introns, exons, 5’UTR, 3’UTR, see **Figure S4d**). ***P= 0.0001 between groups #1 and 2, *P= 0.02 between groups #2 and 3, ^NS^P= 0.1 between groups #3 and 4 (Wilcoxon test).

**Figure S5. a** CpGs in M-ATAC peaks that overlap between both replicates were divided into three groups according to methylation status from M-ATAC: 1/ Low Methylation (<20%, 423379 CpGs), 2/ Intermediate methylation (20-80, 7390 CpGs), 3/ High methylation (>80%, 162 CpGs). **b** CpGs in M-ATAC peaks were divided into three groups according to methylation status from WGBS: 1/ Low Methylation (<20%, 351561 CpGs), 2/ Intermediate methylation (20-80, 58655 CpGs), 3/ High methylation (>80%, 17385 CpGs). Of note, a cutoff of 5 reads coverage were applied, and as opposed to **Fig. 3a**, no additional division was made based on coverage. ***P <0.001 (Wilcoxon text). **c** Transcriptional output for the 4 groups from **Fig. 3a**, for the CpGs in M-ATAC peaks at intragenic regions (introns, exons, 5’UTR, 3’UTR, see **Fig. 3b**). *P= 0.028 between groups #1 and 2, ***P= 1.38e-38 between groups #2 and 3, ^NS^P= 0.88 between groups #3 and 4 (Wilcoxon test). **d** Heatmaps of M-ATAC from individual replicates compared to merged replicates for the 4 groups of CpGs in M-ATAC peaks from **Fig. 3a. e** Heatmaps of M-ATAC, H3K4me1, H3K4me3 and H3K27ac signal corresponding to the average profiles shown in **Fig. 3d** for the 4 groups of CpGs in M-ATAC peaks from **Fig. 3a** at promoters (left panel) versus non-promoters (right panel). **f** Table showing the number of CpGs, M-ATAC and CTCF M-ChIP peaks, CTCF motifs and CpGs within CTCF motifs for the 4 groups of CpGs in M-ATAC peaks from **Fig. 3a**.

**Figure S6**. CTCF M-ChIP. **a** Comparison of CTCF motifs found using CTCF ChIP-seq and CTCF M-ChIP. **b** Scatter plot showing the relationship between CTCF enrichment and CpG methylation within the CTCF motifs in CTCF M-ChIP peaks. **c** CpGs within CTCF motifs in M-ChIP peaks that overlap between both replicates were divided into three groups according to methylation status from M-ChIP: 1/ Low Methylation (<20%, 20644 CpGs), 2/ Intermediate methylation (20-80, 3809 CpGs), 3/ High methylation (>80%, 328 CpGs). **d** Heatmaps of CTCF M-ChIP signal from individual replicates compared to merged replicates for the 3 groups of CpGs in M-ChIP peaks from **Figure S6c** and average profile for merged replicates. **e** Representative IGV screenshots of the 3 groups of CpGs in a CTCF motif within a CTCF peak shown in **Figure S6c** based on methylation levels from CTCF M-ChIP. **f** Table showing the number of CpGs and CTCF M-ChIP peaks depending on methylation levels of either all CpGs in CTCF peaks or only CpGs within a CTCF motif. **g** Scatter plot showing probability of the CpGs in a CTCF motif for **Figure S3c** being in their assigned group.

**Figure S7**. Methylation at C2 and C12 CpGs within CTCF motif. **a** Tables and histogram representing the number of cytosines at position C2 and C12 in the CTCF motif MA0139.1 in CTCF M-ChIP peaks as well as the frequency of the observed versus expected co-occurrence of methylation at C2 and C12 (χ2 = 1531, p-value < 0.001), and the number of CTCF motifs for each C2/C12 methylation combination. **b** Heatmap and average profile of CTCF M-ChIP signal at CTCF motifs with C2/C12 methylation combinations. **c** Frequency of methylation in the CTCF motif from CTCF M-ChIP, for the 7 possible combinations of base variations associated with C at positions 2 (1st couple of nucleotides) and 12 (2d couple of nucleotides).

**Figure S8**. KLF4 mutation. **a** Sequencing from the IRES reverse primer of WT versus Mutant Klf4. The reverse complement of the sequence highlights the wild type (Arginine, R) and mutant (Alanine, A) Klf4. **b** Western blot showing the levels of KLF4 protein and H3 bulk histone in WT mESC, *Kfl4*^*-/-*^ mESC and *Klf4*^*-/-*^ mESC that expressed either a WT or a mutant version of KLF4. **C** Heatmap showing the binding profile of WT and mutant KLF4 R462A duplicates at WT specific (1836), mutant specific (267) and common (303) KLF4 M-ChIP peaks defined in **Fig. 4d. d** Average cytosine methylation from M-ATAC versus KLF4 M-ChIP in WT KLF4 expressing cells (top panel, Pearson Correlation = 0.78, p-value < 2.2e-16) and from KLF4 M-ChIP versus WGBS (bottom panel, Pearson Correlation = 0.84, p-value < 2.2e-16) for CpGs within WT specific KLF4 M-ChIP peaks.

## Availability of data and materials

All raw and processed sequencing data generated in this study have been submitted to the NCBI Gene Expression Omnibus (GEO; http://www.ncbi.nlm.nih.gov/geo/) under accession number GSE129673. The following secure token has been created to allow review of record GSE129673 while it remains in private status: Shkzikwortmhhal.

## Author contributions

P.L and F.I designed the EpiMethylTag experiments. S.V, V.S and D.C.D.G designed and performed the experiments involving KLF4. P.L performed the ChIP/M-ChIP/ATAC/M-ATAC experiments. G.S, M.C and K-T.K developed the analytical pipeline. PL, G.S, F.I, T.S, S.B, M.C and K-T.K analyzed the data. J.S, E.A, M.S and J.S supervised the study. P.L and J.S wrote the paper.

## Ethics approval and consent to participate

Not applicable

## Acknowledgements

The authors thank the New York University School of Medicine High Performance Computing (HPC) for computing technical support, and Adriana Heguy and the Genome Technology Center (GTC) core for sequencing efforts.

