## Supplementary figures for "EpiMethylTag simultaneously detects ATAC-seq or ChIP-seq signals with DNA methylation"

### Figure S1

a Number of peaks per sample

| | peaks # |
|---|---|
| M-ATAC-rep1 | 31907 |
| M-ATAC-rep2 | 30413 |
| ATAC-rep1 | 28717 |
| ATAC-rep2 | 37537 |
| M-ATAC-NC-rep1 | 31806 |
| M-ATAC-NC-rep2 | 36710 |
| CTCF M-ChIP-rep2 | 33898 |
| CTCF M-ChIP-rep1 | 32195 |
| CTCF ChIP-rep1 | 41731 |
| CTCF ChIP-rep2 | 52327 |
| CTCF M-ChIP-NC-rep1 | 22576 |
| CTCF M-ChIP-NC-rep2 | 31409 |

b Jaccard indexes of peaks intersections between samples

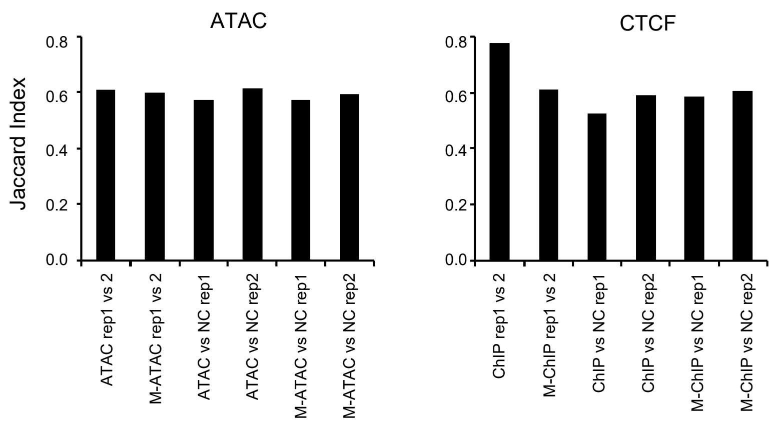


c Correlation of reads counts within peaks

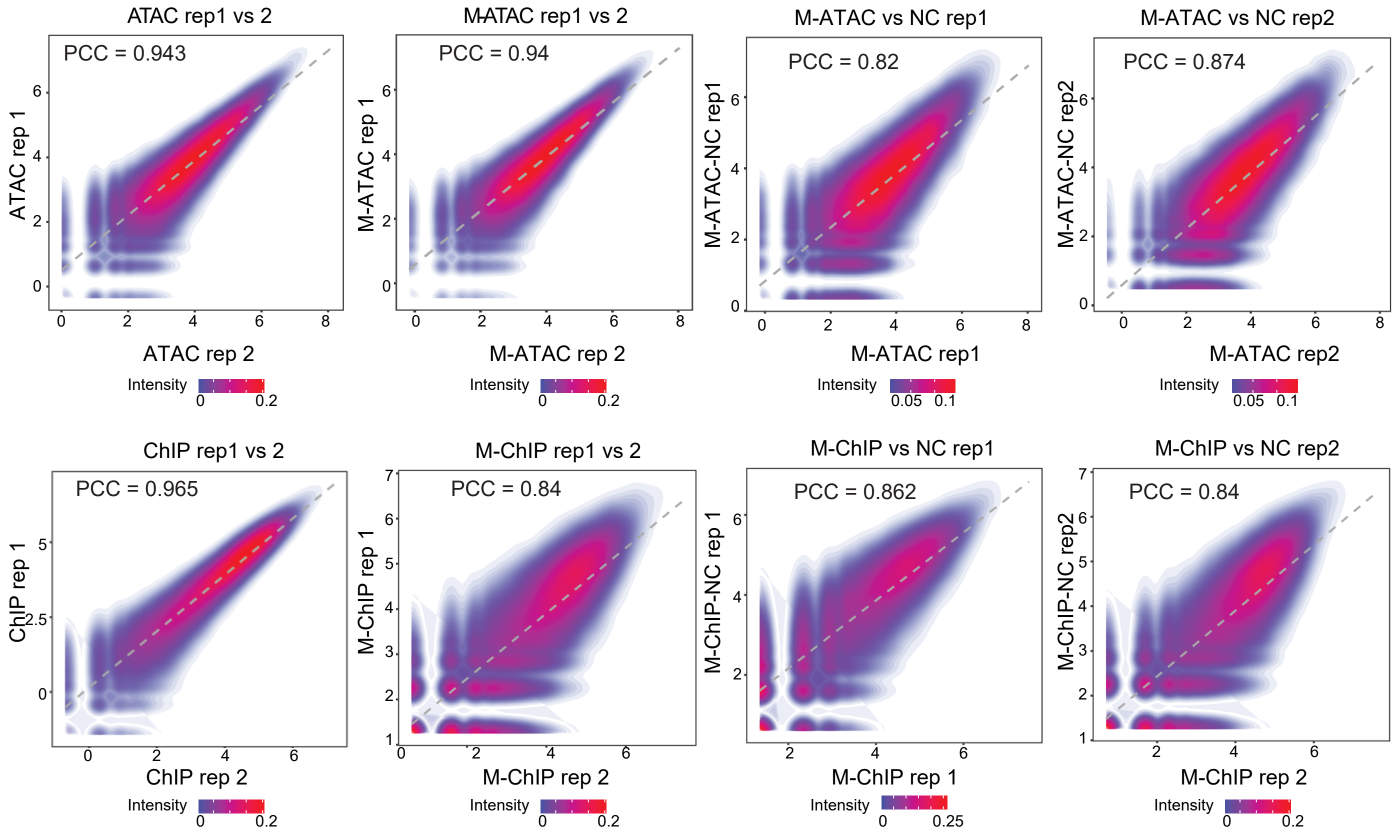

Figure S2

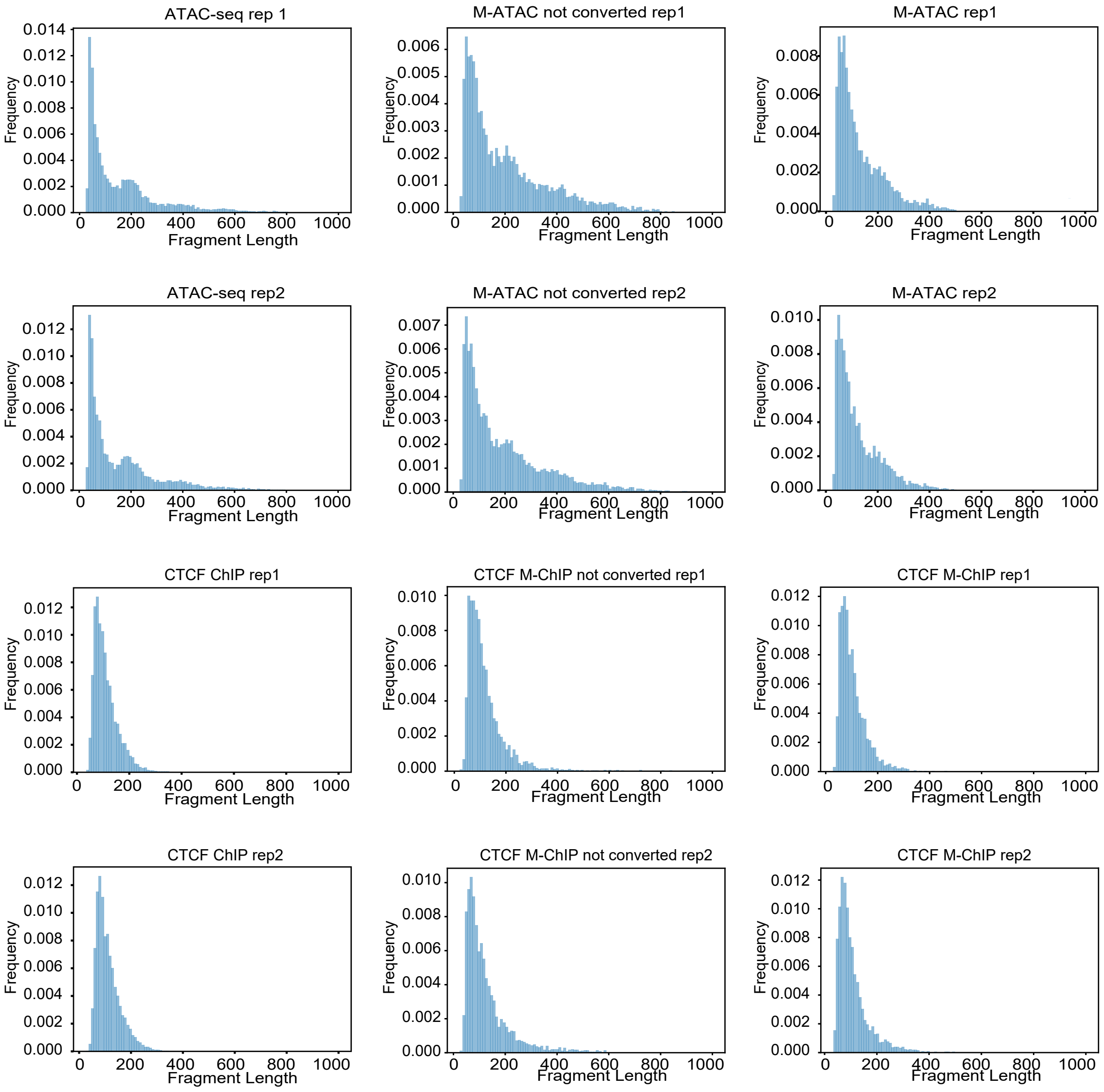
ATAC-seq rep 1
M-ATAC not converted rep1
M-ATAC rep1
ATAC-seq rep2
M-ATAC not converted rep2
M-ATAC rep2
CTCF ChIP rep1
CTCF M-ChIP not converted rep1
CTCF M-ChIP rep1
CTCF ChIP rep2
CTCF M-ChIP not converted rep2
CTCF M-ChIP rep2
Frequency
Fragment Length

### Figure S3

Average cytosine methylation relative to its position in M-ATAC peak

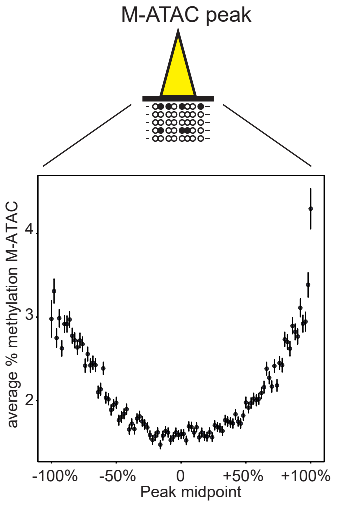

### Figure S4

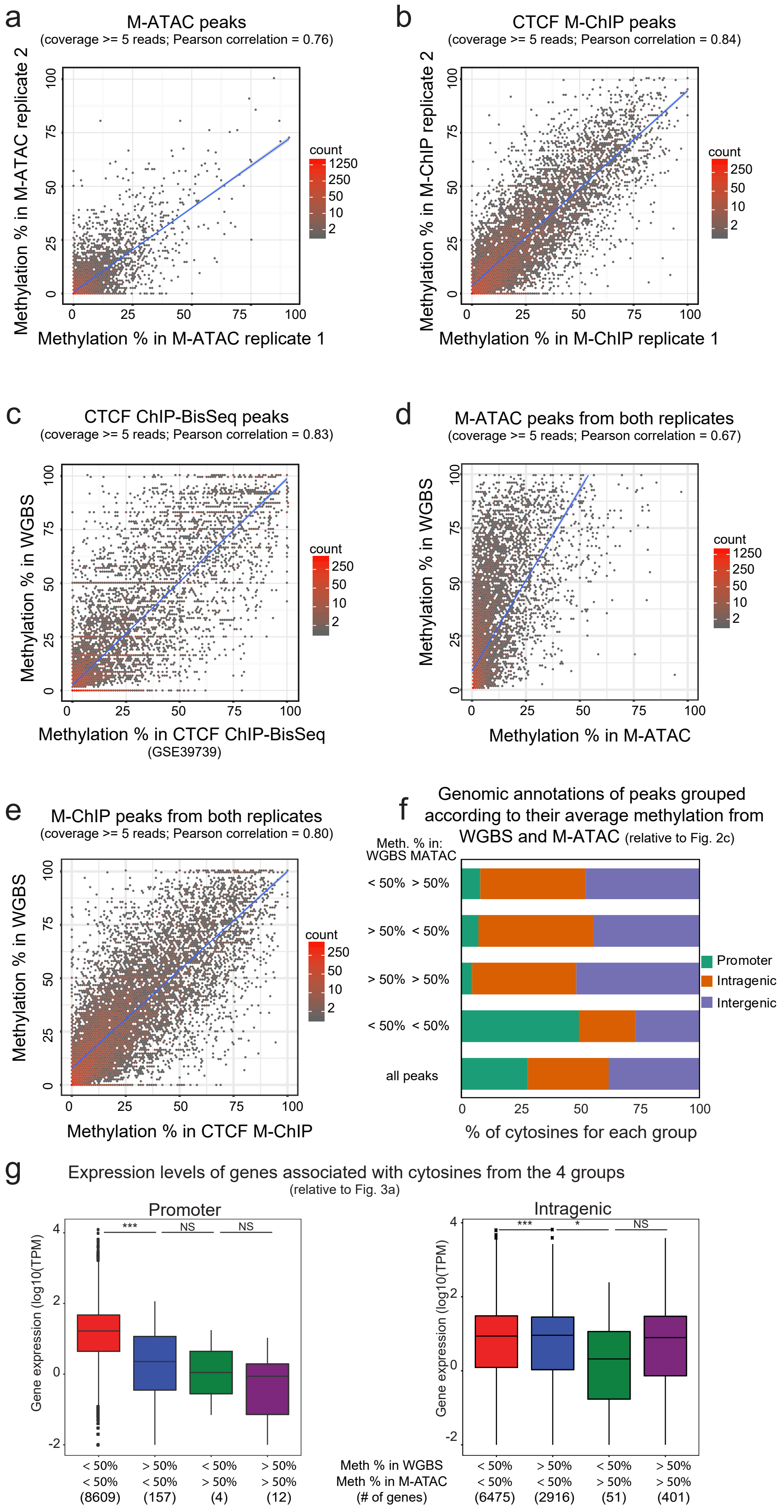

a
M-ATAC peaks
(coverage >= 5 reads; Pearson correlation = 0.76)
Methylation % in M-ATAC replicate 2
Methylation % in M-ATAC replicate 1
count
1250
250
50
10
2
b
CTCF M-ChIP peaks
(coverage >= 5 reads; Pearson correlation = 0.84)
Methylation % in M-ChIP replicate 2
Methylation % in M-ChIP replicate 1
c
CTCF ChIP-BisSeq peaks
(coverage >= 5 reads; Pearson correlation = 0.83)
Methylation % in WGBS
Methylation % in CTCF ChIP-BisSeq
(GSE39739)
d
M-ATAC peaks from both replicates
(coverage >= 5 reads; Pearson correlation = 0.67)
Methylation % in WGBS
Methylation % in M-ATAC
e
M-ChIP peaks from both replicates
(coverage >= 5 reads; Pearson correlation = 0.80)
Methylation % in WGBS
Methylation % in CTCF M-ChIP
f
Genomic annotations of peaks grouped according to their average methylation from WGBS and M-ATAC (relative to Fig. 2c)
Meth. % in:
WGBS MATAC
< 50% > 50%
> 50% < 50%
> 50% > 50%
< 50% < 50%
all peaks
Promoter
Intragenic
Intergenic
% of cytosines for each group
g
Expression levels of genes associated with cytosines from the 4 groups
(relative to Fig. 3a)
Promoter
Intragenic
Gene expression (log10(TPM))
***
NS
*
Meth % in WGBS
Meth % in M-ATAC
(# of genes)
< 50% < 50% (8609)
> 50% < 50% (157)
< 50% > 50% (4)
> 50% > 50% (12)
< 50% < 50% (6475)
> 50% < 50% (2916)
< 50% > 50% (51)
> 50% > 50% (401)

### Figure S5

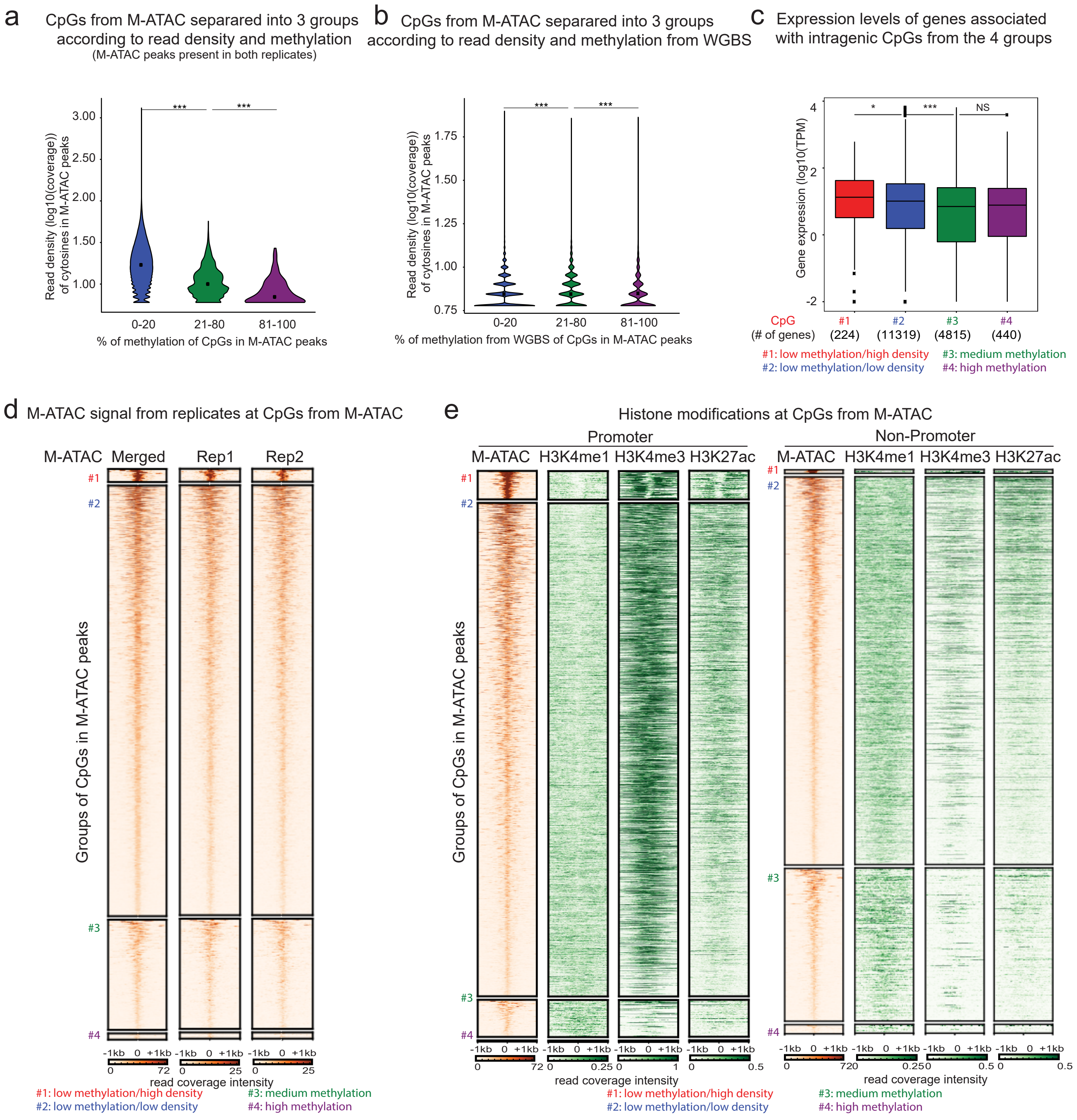


f CpGs from M-ATAC separared into 4 groups according to read density and methylation (M-ATAC peaks from merged replicates)

| M-ATAC group (Fig. 3a) | 1 | 2 | 3 | 4 |
|---|---|---|---|---|
| number CpGs | 22932 | 1348931 | 39321 | 1652 |
| number M-ATAC peaks | 1858 | 44651 | 17069 | 1137 |
| number CTCF M-ChIP peaks | 122 | 6526 | 1845 | 80 |
| number CTCF motifs | 206 | 11062 | 734 | 28 |
| number unique CpGs within CTCF motifs (used for Fig. 4a) | 288 | 17133 | 758 | 25 |
| percentage of CpGs within CTCF motifs | 1.26 | 1.27 | 1.93 | 1.51 |

### Figure S6

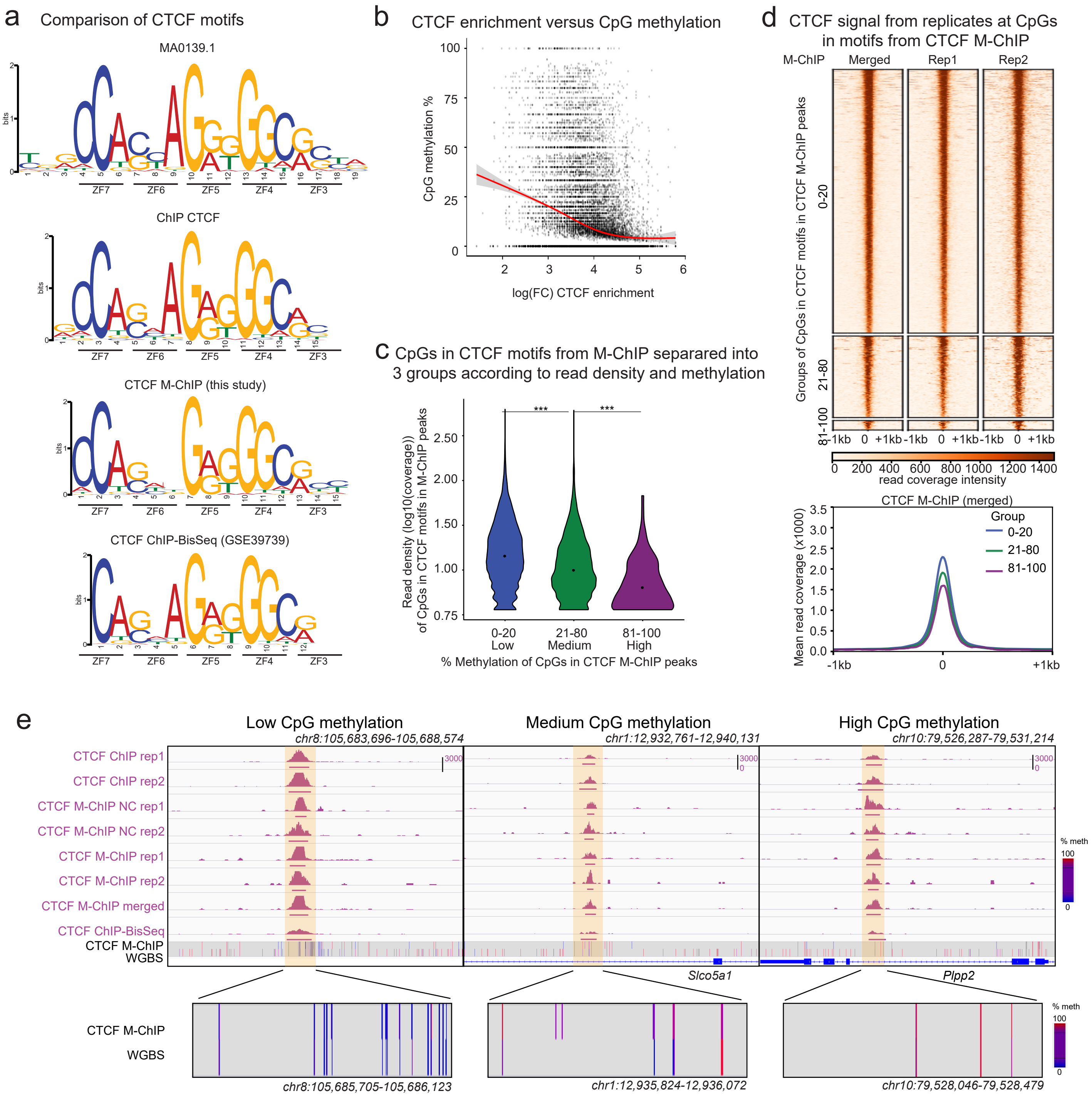


f

Number of CpGs and CTCF M-ChIP peaks in the 3 groups of CpGs based on methylation levels from M-ChIP

| | CpGs within a CTCF peak | | | CpGs in a CTCF motif within a peak | | |
|---|---|---|---|---|---|---|
| | 0-20 Low | 21-80 Medium | 81-100 High | 0-20 Low | 21-80 Medium | 81-100 High |
| number of CpGs (no cutoff) | 103302 | 40389 | 6397 | 23524 | 4949 | 551 |
| number of CpGs (cutoff 5) | 81443 | 28793 | 3359 | 20644 | 3809 | 328 |
| number of peaks (cutoff 5) | 14460 | 12272 | 2347 | 8252 | 2551 | 296 |

g

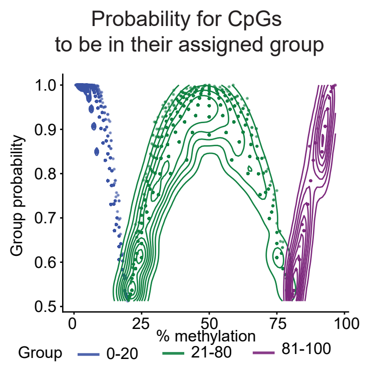

### Figure S7

**a** C2/C12 methylation in CTCF motif at single molecules of CTCF M-ChIP DNA

| Observed | | C2 | % | C12 | % |
|---|---|---|---|---|---|
| Unmethylated | ○ | 8829 | 97.77 | 8817 | 97.64 |
| Methylated | ● | 201 | 2.23 | 213 | 2.36 |

| C2 | C12 | Observed reads # | Observed reads % | Expected reads % | Motifs # |
|---|---|---|---|---|---|
| ○ | ○ | 8704 | 96.39 | (97.77 * 97.64)/100 = 95.47 | 1320 |
| ○ | ● | 125 | 1.38 | (97.77 * 2.36)/100 = 2.31 | 113 |
| ● | ○ | 113 | 1.25 | (2.23 * 97.64)/100 = 2.17 | 89 |
| ● | ● | 88 | 0.97 | (2.23 * 2.36)/100 = 0.053 | 58 |
| | | 9030 | | | |

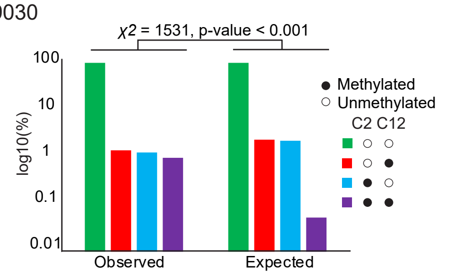


**b** CTCF binding signal at motifs with C2/C12 methylation combinations

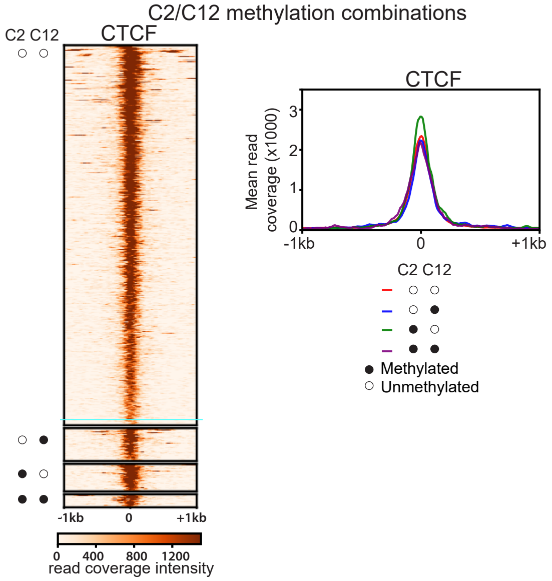


**c** Methylation frequency for the different combination of C2 and C12 di-nucleotides

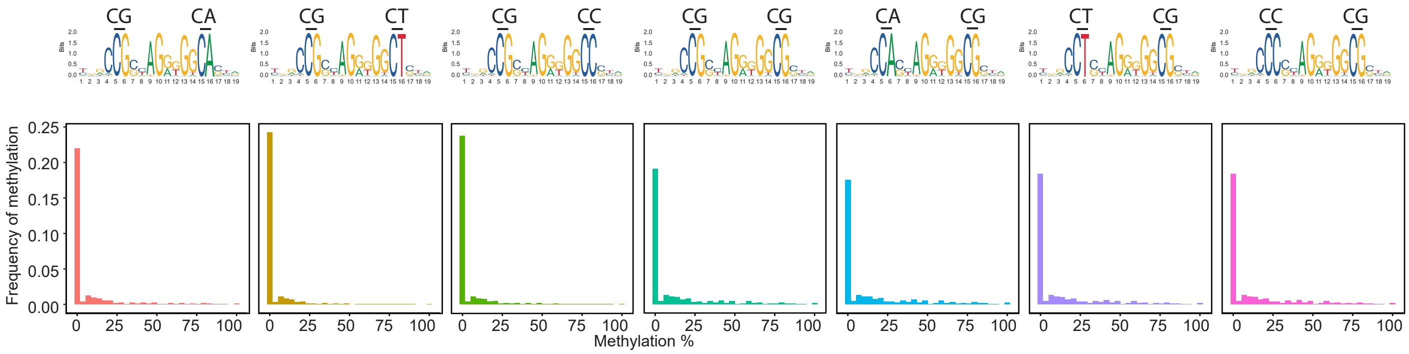

Figure S8

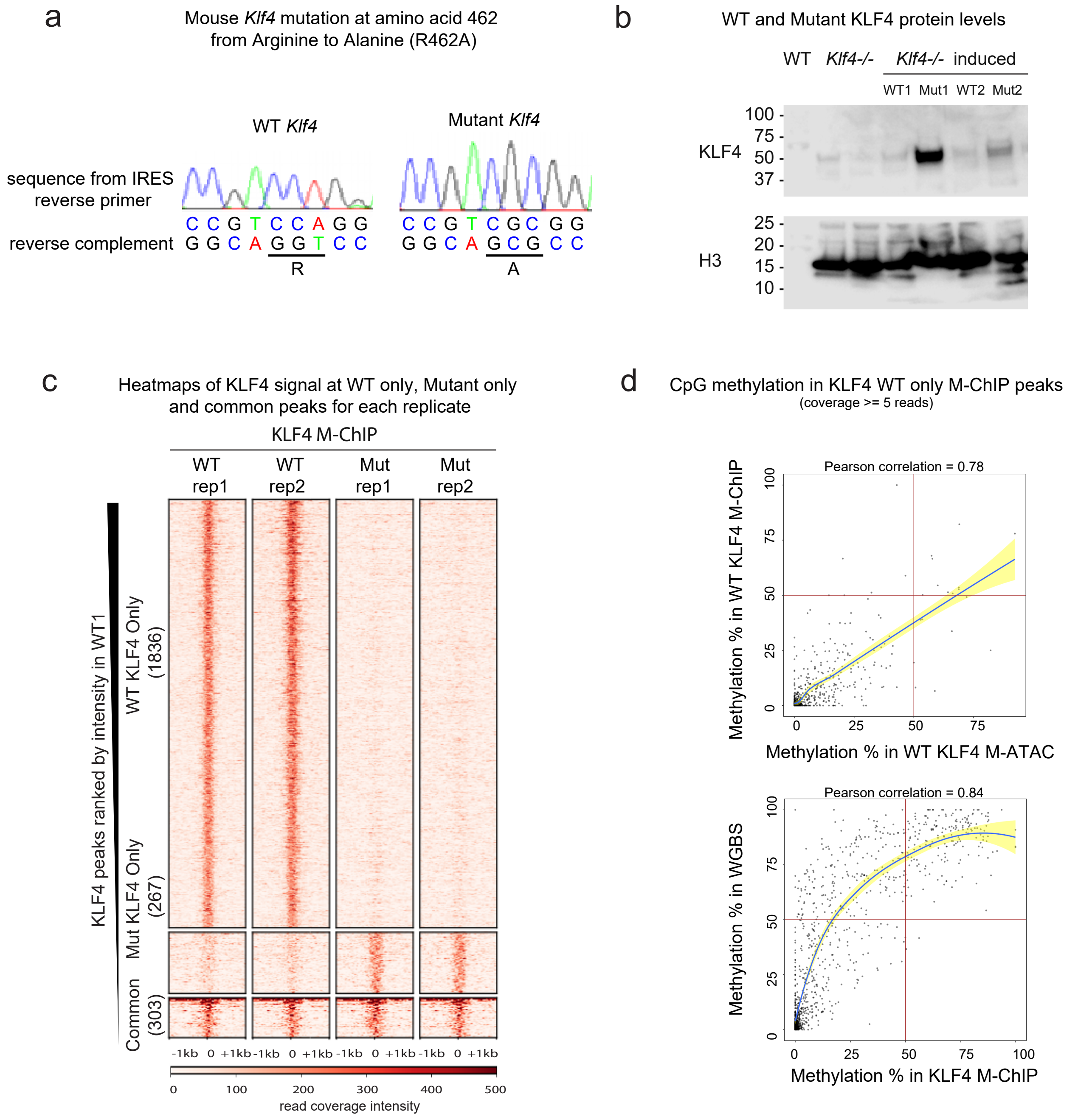
a
Mouse Klf4 mutation at amino acid 462 from Arginine to Alanine (R462A)
WT Klf4
Mutant Klf4
sequence from IRES reverse primer
reverse complement
C C G T C C A G G
G G C A G G T C C
R
C C G T C G C G G
G G C A G C G C C
A
b
WT and Mutant KLF4 protein levels
WT
Klf4-/-
Klf4-/- induced
WT1 Mut1 WT2 Mut2
KLF4
100
75
50
37
H3
25
20
15
10
c
Heatmaps of KLF4 signal at WT only, Mutant only and common peaks for each replicate
KLF4 M-ChIP
WT rep1
WT rep2
Mut rep1
Mut rep2
KLF4 peaks ranked by intensity in WT1
WT KLF4 Only (1836)
Mut KLF4 Only (267)
Common (303)
-1kb 0 +1kb
0 100 200 300 400 500
read coverage intensity
d
CpG methylation in KLF4 WT only M-ChIP peaks
(coverage >= 5 reads)
Pearson correlation = 0.78
Methylation % in WT KLF4 M-ChIP
Methylation % in WT KLF4 M-ATAC
Pearson correlation = 0.84
Methylation % in WGBS
Methylation % in KLF4 M-ChIP
0 25 50 75 100
